# Comparison of mitochondrial response to SARS-CoV-2 spike protein receptor binding domain in human lung microvascular, coronary artery endothelial and bronchial epithelial cells

**DOI:** 10.1101/2023.12.19.572363

**Authors:** Gabrielė Kulkovienė, Deimantė Narauskaitė, Agilė Tunaitytė, Augusta Volkevičiūtė, Zbigniev Balion, Olena Kutakh, Dovydas Gečys, Milda Kairytė, Martyna Uldukytė, Edgaras Stankevičius, Aistė Jekabsone

## Abstract

Recent evidence indicate that SARS-CoV-2 spike protein affects mitochondria with a cell type-dependent outcome. We elucidate the effect of SARS-CoV-2 receptor binding domain (RBD) on the mitochondrial network and cristae morphology, oxygen consumption, mitoROS production, and inflammatory cytokine expression in cultured human lung microvascular (HLMVEC) and coronary artery endothelial (HCAEC) and bronchial epithelial cells (HBEC). Live Mito Orange staining, STED microscopy and Fiji MiNa analysis were used for mitochondrial cristae and network morphometry, Agilent XFp analyser for mitochondrial/glycolytic activity, MitoSOX fluorescence for mitochondrial ROS, and qRT-PCR plus Luminex for cytokines. In HLMVEC, SARS-CoV-2 RBD fragmented the mitochondrial network, decreased cristae density, mitochondrial oxygen consumption and glycolysis and induced mitoROS-mediated GM-CSF and IL-1β expression in all three investigated cell types and IL-8 - in both endothelial cell types. Mitochondrial ROS control SARS-CoV-2 RBD-induced inflammation in HLMVEC, HCAEC and HBEC, with the mitochondria of HLMVEC being more sensitive to SARS-CoV-2 RBD.

## 1. Introduction

COVID-19 causes severe pulmonary and cardiovascular complications via mitochondrial signalling and immuno-metabolic reprogramming-mediated inflammation [1]. SARS-CoV-2 infection disrupts the mitochondrial network, alters respiratory chain function and increases mitochondrial reactive oxygen species (mitoROS) [2–4], leading to oxidative stress in severe cases of COVID-19 [5]. This study compares the involvement of mitochondria and mitoROS in the inflammation triggered by the SARS-CoV-2 spike glycoprotein receptor binding domain (SCoV2-RBD) in human lung microvascular and coronary artery endothelial, and bronchial epithelial cells, with a specific focus on the mitochondrial network integrity and cristae density.

## 2. Results

### 2.1 The effect of SCoV2-RBD on mitochondrial morphology

First, we evaluated changes in mitochondrial morphology of human lung microvascular endothelial (HLMVEC), coronary artery endothelial (HCAEC) and bronchial epithelial cells (HBEC) after 24-hour SCoV2-RBD treatment. Visual examination of the mitochondrial network revealed higher fragmentation in SCoV2-RBD-affected HLMVEC (Fig. 1a). Treatment with SCoV2-RBD decreased mitochondrial area, or footprint, by 25%, length of branches by 11.6%, summed branch length by 20.9%, and number of network branches by 25.9% compared to healthy cells (Fig. 1b). There were no visual qualitative of significant quantitative changes in HCAEC and HBEC after SCoV2-RBD treatment (Fig. A3, A4).

**Figure 1.**
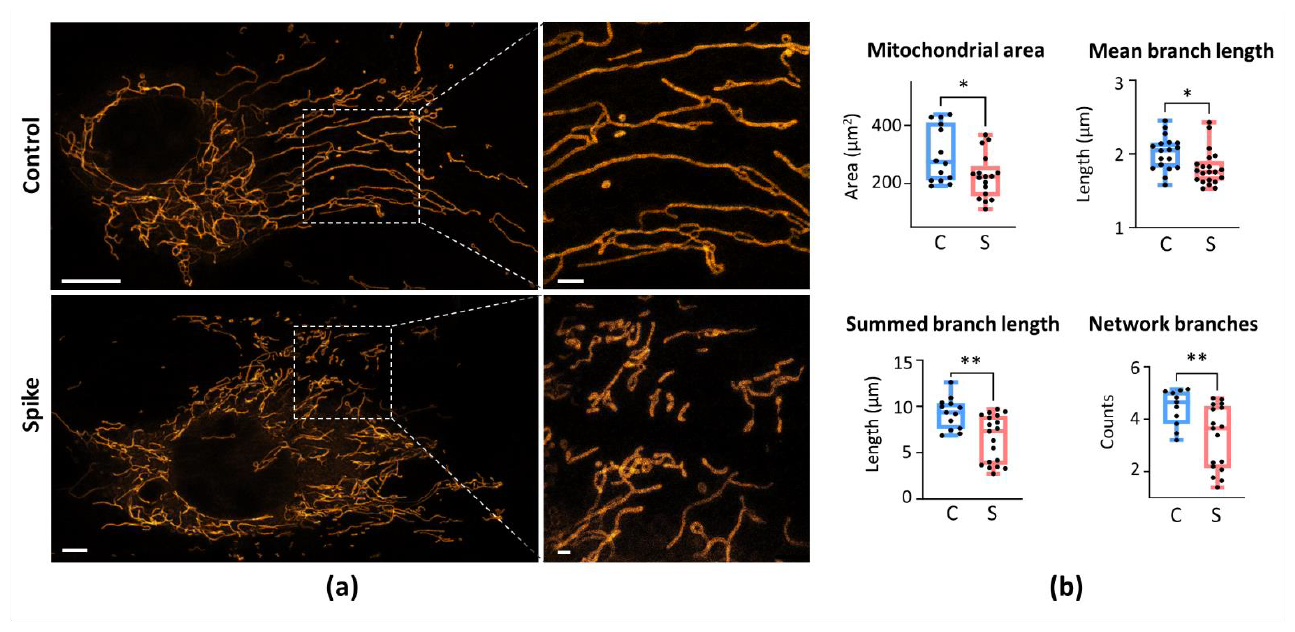
The effect of SCoV2-RBD on mitochondrial network morphology of HLMVEC. In (a) - representative images of mitochondrial network of untreated (Control) and SCoV2-RBD-treated HLMVEC (Spike). Live Mito Orange and images stained the cells were taken by an Olympus IX83 confocal microscope equipped with a STEDYCON STED nanoscope. The scale bar is 5 μm for the mitochondrial network of entire cells and 1 μm for the ROIs. In (b) – the quantitative results of mitochondrial network parameters calculated by means of the MiNa plugin of Fiji software. Results are normalised to untreated cells (Control) and represented as fold changes. C is for Control, S – for Spike, which means SCoV2-RBD treatment. ^*^p<0.05, ^**^p < 0.01; one-way ANOVA Tukey’s test

Moreover, the intercristae distance in SCoV2-RBD-primed HLMVEC increased significantly by 30% compared to untreated cells (Fig. 2). However, no structural alterations in mitochondrial cristae were observed in HCAEC and HBEC (data not shown).

**Figure 2.**
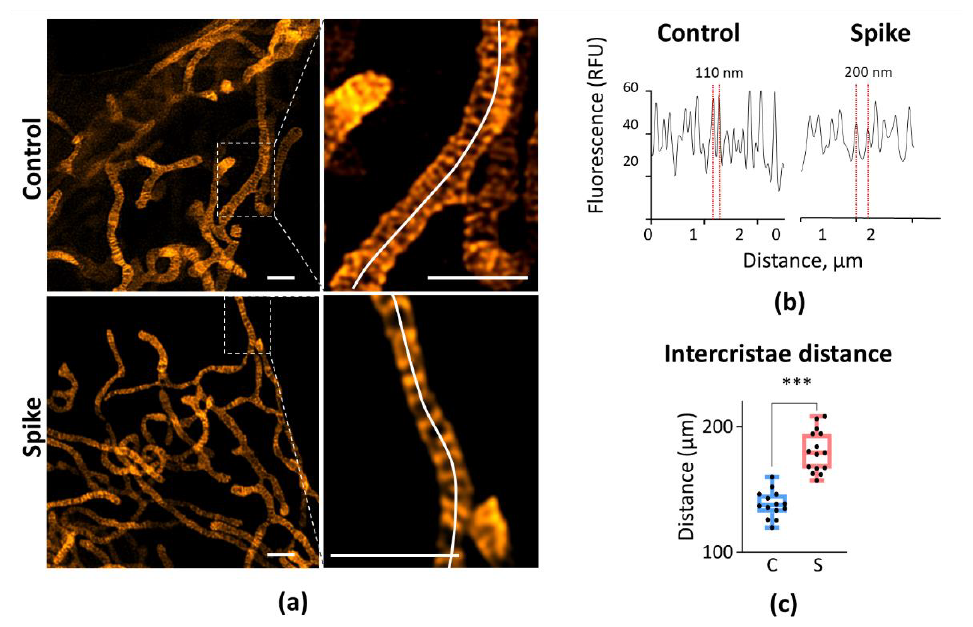
The effect of 24h treatment of SCoV2-RBD on HLMVEC cristae density. In (**a**) – representative images of untreated (Control) and SCoV2-RBD-treated (Spike) cells. The dotted line in the images on the left indicates regions of interest presented on the right. The scale bar is 1 μm. In (b) – the distance between mitochondrial cristae evaluated according to the fluorescence intensity measured along the curved white line on the mitochondrial network branch in (**a**), right. In (**c**) – the quantitative evaluation of intercristae distance in control (C) and SCoV2-RBD-treated (S) cells. ^***^p < 0.001; one way ANOVA Tukey’s test.

### 2.2 The effect of SCoV2-RBD on mitochondrial and glycolytic activity and mitoROS

Further, mitochondrial respiration and glycolysis activity were evaluated after priming HLMVEC, HCAEC, and HBEC cultures with SCoV2-RBD. The mito-morphological events in HLMVEC were accompanied by the significant suppression of mitochondrial respiration and glycolysis (Fig. 3). A significant decrease was observed in basal and maximal oxygen consumption and glycolysis. However, any substantial changes in the energetic activity of HCAEC and HBEC were determined. These results suggest that SCoV2-RBD impairs mitochondrial function in HLMVEC cells by disrupting their network and cristae morphometrics.

**Figure 3.**
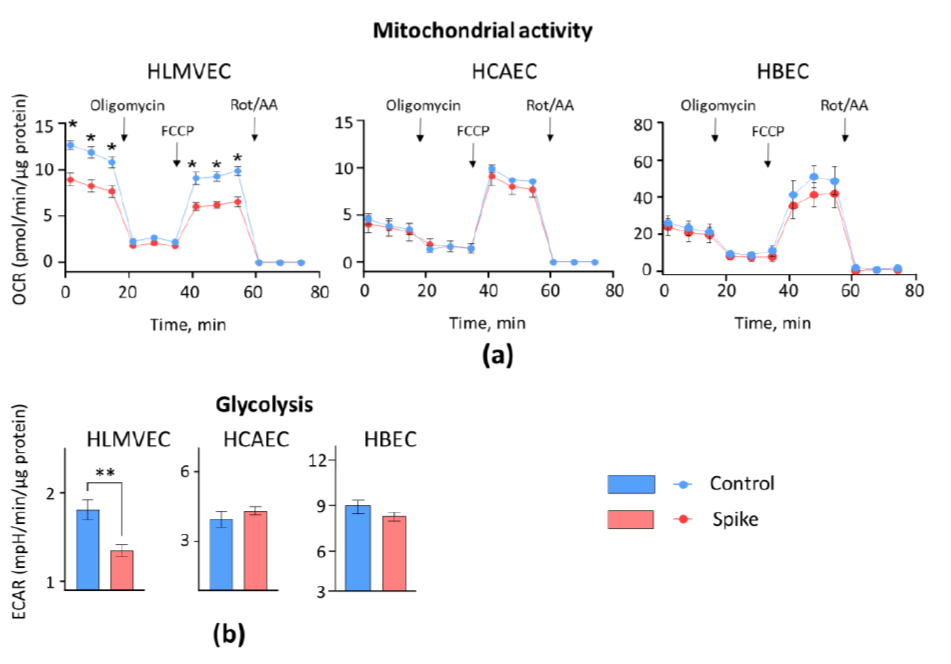
The effect of 24h treatment of SCoV2-RBD on mitochondrial (**a**) and glycolytic (**b**) activity of HLMVEC, HCAEC and HBEC. In (**a**), oxygen consumption rate (OCR) represents the efficiency of mitochondrial respiration. The initial three measurements indicate the basal level of mitochondrial respiration. The next three measurements, taken after introducing oligomycin to inhibit ATP synthase, show the oxygen consumption caused by proton leak. After that, another three measurements represent the maximum capacity of mitochondrial oxygen consumption, which happens when FCCP uncouples the inner mitochondrial membrane. Lastly, the last three measurements account for oxygen consumption unrelated to mitochondria, arising when the mitochondrial respiratory chain is hampered by rotenone and antimycin A. In (b), the extracellular medium acidification rate (ECAR) represents the efficacy of glycolysis. ^*^p<0.05, ^**^p < 0.01; one way ANOVA Tukey’s test.

Changes in mitochondrial respiratory chain activity might affect ROS production. Therefore, further in this study, the effect of SCoV2-RBD protein on mitoROS generation in HLMVEC, HCAEC and HBEC was investigated. The primary ROS produced within mitochondria is a superoxide, which is incapable of crossing membranes and stays at the production site. A fluorescent probe targeting superoxide in mitochondria called MitoSOX red was employed to measure it. SCoV2-RBD significantly induced mitoROS production in HLMVEC and HCAEC cells: MitoSOX fluorescence intensity increased by 2.6% and 7.4 %, respectively, compared to untreated cells (Fig. 4). No change of mitoROS generation in HBEC cells was detected.

**Figure 4.**
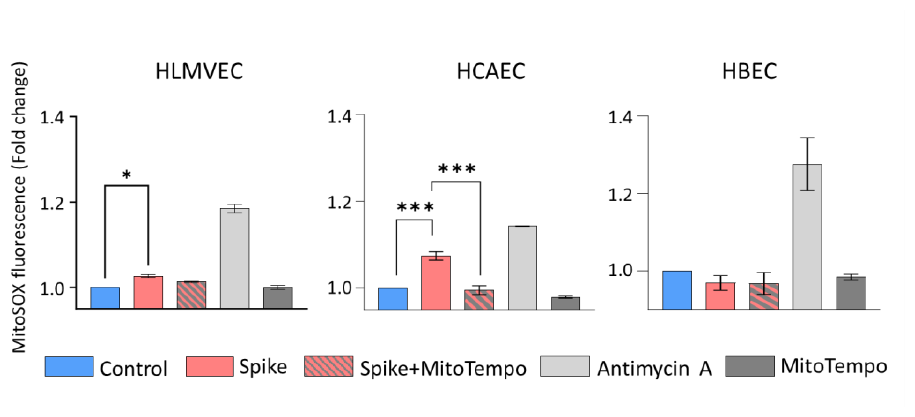
MitoROS production by HLMVEC, HCAEC and HBEC after SCoV2-RBD treatment. Spike - SCoV2-RBD, MitoTempo – specific mitochondrial superoxide scavenger used to test assay specificity. Antimycin A - a positive control. Results are normalised to untreated cells (Control) and represented as fold changes. ^*^p<0.05, ^***^p < 0.001; one way ANOVA Tukey’s test.

### 2.3 The impact of mitoROS on SCoV2-RBD-induced effect on mitochondrial morphology

MitoROS production might result from mitochondrial network morphology changes, but it also might be the cause of such changes. We have applied a specific mitochondrial superoxide scavenger, MitoTempo, to test whether mitoROS influenced the mitochondrial network changes that occurred after SCoV2-RBD treatment in HLMVEC cells. Interestingly, MitoTempo was able to rescue mitochondrial fragmentation in HLMVEC (Fig. 5a). It restored the summed branch length and number of network branches’ values to the basal level (Fig. 5b), indicating that mitochondrial elongation and interconnectivity were back to the untreated control level. However, mitoROS had no detectable impact on SCoV2-RBD effect on mitochondrial footprint area and mean branch length (Fig. A5), indicating that mitoROS were not in control of SCoV2-RBD-induced mitochondrial loss.

**Figure 5.**
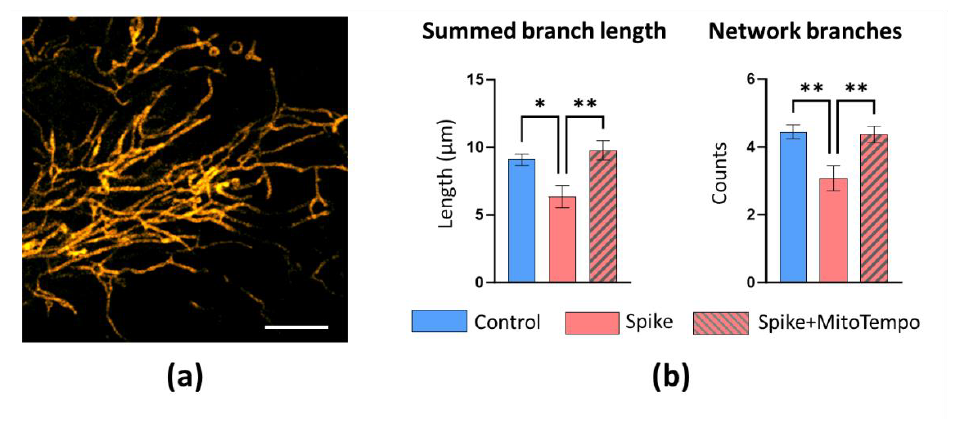
The impact of mitoROS in SCoV2-RBD-induced mitochondrial network alteration in HMLVEC. Spike - SCoV2-RBD. In (**a**) – a representative image of the restored HLMVEC mitochondrial network after MitoTempo addition. In (**b**) – column charts represent MitoTempo effects on quantitative mitochondrial network parameters. Results are normalised to untreated cells (Control) and represented as fold changes. ^*^p<0.05, ^**^p < 0.01; one way ANOVA Tukey’s test.

### 2.4 The impact of mitoROS on SCoV2-RBD-induced inflammatory cytokines

MitoROS are involved in multiple cell signalling pathways, including inflammation. Thus, further in the study, we have tested whether inflammatory cytokine expression and secretion after SCoV2-RBD treatment is affected by mitoROS scavenger mitoTempo in HLMVEC, HCAEC and HBEC.

SCoV2-RBD treatment resulted in significant cytokine upregulation in all cell lines when compared to untreated cells (Fig. 6). In HLMVEC, NF-kβ expression increased by 1.15 fold, GM-CSF – even 4.25, IL-8 - 1.5, and IL-1β - 3.1. In HCAEC, GM-CSF expression was increased by 3.5 fold, IL-8 by 1.35, IL-1β by 1.9, and IL-6 by 1.3. In HBEC, average NF-kβ expression level was increased by 1.2 fold, CCL2 by 2.3, GM-CSF – by 5.1, IL-8 by 1.7, and IL-1β by 4.6.

**Figure 6.**
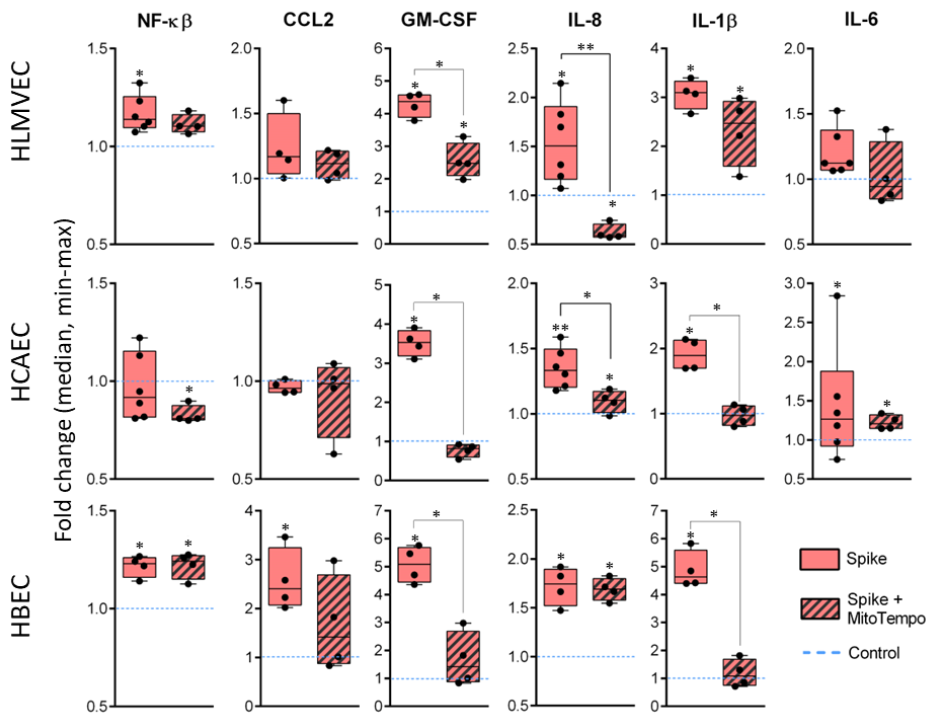
NF-kβ, CCL2, GM-CSF, IL-8, IL-1β and IL-6 gene expression changes in HLMVEC, HCAEC and HBEC3 cells after treatment with of SCoV2-RBD (Spike). Treatment resulted in significant cytokine-coding gene expression upregulation across all cell lines. The addition of MitoTempo (Spike+MitoTempo) reversed gene expression changes caused by SCoV2-RBD in GM-CSF, IL-8 and IL-1β. Results are normalised to untreated cells (Control) and represented as fold change values. ^*^p<0.05; ^**^p<0.01; Mann-Whitney U test.

The addition of MitoTempo during SCoV2-RBD treatment significantly prevented SCoV2-RBD-induced gene upregulation down to the baseline value or even induced downregulation. Significant differences between SCoV2-RBD and MitoTempo-supplemented SCoV2-RBD treatment were observed in GM-CSF (fold changes, or FC= 4.25 vs. 2.5) and IL-8 (FC= 1.5 vs. 0.6) in HLMVEC; GM-CSF (FC= 3.5 vs. 0.9), IL-8 (FC= 1.35 vs. 1.2) and IL-1β (FC= 1.9 vs. 1) in HCAEC; GM-CSF (FC= 5.1 vs. 1.4) and IL-1β (FC= 4.6 vs. 1) expression in HBEC3 cells. Noteworthy, the addition of MitoTempo significantly downregulated expression of IL-8 (FC=0.6) in HLMVEC cells and NF-kβ (FC=0.8) in HCAEC cells. Despite cytokine expression-attenuating effects, GM-CSF and IL-1β remained significantly upregulated in HLMVEC cells. Similar results were observed in IL-8 and IL-6 expression in HCAEC cells as well as IL-8 and IL-1β expression in HBEC3 cells.

SCoV2-RBD treatment significantly increased GM-CSF secretion from HLMVEC and HCAEC cells to their culture medium in a MitoTempo-sensitive manner, clearly indicating involvement of mitoROS in GM-CSF pathway (Fig. 7). In contrast, SCoV2-RBD did not change the secretion of other investigated cytokines and even decreased the secreted level of TNF-α in HLMVEC (Fig. A6). Besides, the levels of IL-1β, IL-6 and TNF-α were below detection limits in both control and SCoV2-RBD-treated samples of HBEC (Fig. A6).

**Figure 7.**
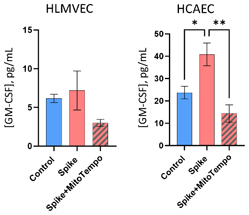
GM-CSF proinflammatory cytokine secretion levels in HLMVEC and HCAEC cell culture medium after SCoV2-RBD alone (Spike) treatment and supplemented with MitoTempo (Spike+MitoTempo). ^*^p<0.05; ^**^p<0.01; one way ANOVA Fisher LSD test.

## 3. Discussion

The study revealed the exclusive sensitivity of HLMVEC mitochondria to SCoV2-RBD signalling. Treatment with SCoV2-RBD protein resulted in a fragmented mitochondrial network (shorter, less branched structures) and decreased mitochondria-covered area, indicating loss of total mitochondrial volume. Moreover, the remaining mitochondria after SCoV2-RBD treatment had larger intercristae space accompanied by suppressed basal and uncoupler-stimulated mitochondrial respiration, pointing to inhibition of the mitochondrial respiratory chain. Similar mitochondrial impairment and fragmentation by SARS-CoV-2 subunit 1 (which also contains the receptor binding domain) was recently observed in primary human ventricular cardiomyocytes [6]. Additionally, mitochondrial respiratory chain dysfunction was followed by Drp1-mediated fission and AIF-induced apoptosis in Calu-3 (human lung adenocarcinoma-derived airway epithelial) cells after SARS-CoV-2 infection [7]. However, no other studies have been published so far evaluating the effect of the SARS-CoV-2 virus or its components on mitochondrial cristae density. Mitochondrial respiratory efficiency relies on assembling and maintaining mitochondrial respiratory supercomplexes, stabilising the individual complex structure and their performance by substrate channelling [8]. The best-known mitochondrial supercomplexes are respirasomes composed of complexes I, III and IV [9]. Mitochondrial capacity to keep supercomplexes assembled directly depends on cristae shape, namely, the tightness of cristae invaginations and the distance between the cristae-forming membranes [10]. Such findings suggest cell interaction with SCoV2-RBD affects mitochondrial respiratory chain complex efficiency, destabilising them by decreasing cristae compactness and making them less dense. OPA1 interaction with Drp1 is shown to be crucial in the formation of tight, supercomplexes-supporting mitochondrial cristae shape [10,11]. Although Drp1 involvement in SARS-CoV-2-induced mitochondrial deficiency is demonstrated in Calu-3 cells [7], no data about Drp1 and OPA1 regulation by SARS-CoV-2 virus or its proteins have been published so far, indicating the signalling sequence from ACE2 receptor binding to mitochondrial cristae reshaping yet has to be discovered. Besides, other than cristae shaping mechanisms of SARS-CoV-2-mediated mitochondrial dysfunction, such as direct downregulation of respiratory chain complex I and ATP synthase genes, as reported by Archer and colleagues [7], might also occur.

Despite HLMVEC mitochondria showing dramatic changes over 24 hours with SCoV2-RBD, no network, cristae or activity changes were detected in HCAEC and HBEC cells after the same treatment. Although this might seem controversial in the context of numerous reports about SARS-CoV-2-induced mitochondrial damage in vascular endothelial and airway epithelial cells, the explanation might be found in differences between the experimental models, including cell types, mimicking infection (from full virus to spike protein, its subunits, domains or epitopes), treatment duration and concentration. When spike protein and its domains mostly refer to mitochondrial signalling via ACE2 [3][6], viral infection activates mitochondrial immune response via recruitment of mitochondrial antiviral signalling protein MAVS, leading to inhibition of mitochondrial fission and mitochondrial network elongation [12]. Nevertheless, our study indicates that certain cell types, including lung capillary endothelial cells, are more sensitive to SCoV2-RBD signalling to their mitochondria, and this sensitivity might contribute to SARS-CoV-2 (and, potentially, other similar viruses)-induced alveolar damage leading to hypoxemia.

While SARS-CoV-2 was described to induce mitochondrial to glycolytic metabolic shift in endothelial cells [3,13], here, we observed a slow-down in both glycolysis and mitochondrial respiration in HLMVEC, indicating that the cells are switched to low-energy profile. Cell viability (tested by PrestoBlue metabolic activity assay) of all three cell types after 24-hour treatment with SCoV2-RBD was unchanged (data not shown); thus, cell loss did not influence the results. Although COVID-19 infection is reported to increase glycolysis via metabolic changes similar to the Warburg effect in cancerous cells [14], ACE2 signalling has opposite signalling, leading to downregulation of glycolytic enzymes and glycolysis slow-down [15] and suggesting the same or similar mechanism might take place in HLMVEC under SCoV2-RBD treatment. However, glycolytic efficiency was unchanged in HCAEC and HBEC after 24-hour treatment with SCoV2-RBD, indicating that such glycolysis suppression is not the case in bronchial epithelial and coronary endothelial cells.

Lower cristae density increases cristae volume, directly correlating to mitoROS production intensity [16]. Indeed, in HLMVEC cells, SCoV2-RBD-induced mitochondrial suppression was accompanied by an increase in mitoROS. Moreover, mitochondrial network fragmentation (but not mitochondrial degradation) was even mediated by mitoROS. Interestingly, despite no detectable mitochondrial functional or structural changes, an even more prominent mitoROS increase was induced in HCAEC. There was no mitoROS increase in HBEC after 24 hours with SCoV2-RBD; however, this does not exclude the possibility that mitoROS were generated more intensively during some treatment periods before coming back to the control level. ACE2 binding can trigger mitoROS via NOX4 [17]; thus, the descending mitoROS induction in the row of HCAEC, HLMVEC, and HBEC can at least partly be explained by higher expression of ACE2 in cardiovascular tissues compared to the pulmonary and in endothelium compared to the epithelium [18,19].

MitoROS scavenger MitoTempo prevented SCoV2-RBD-induced expression of GM-CSF and IL-1β, in all three investigated cell types and IL-8 in HLMVEC and HCAEC, indicating mitoROS signalling as a key regulator of the genes after RBD-receptor interaction. Moreover, mito-ROS-dependent induction of GM-CSF expression was confirmed by the secretion of this cytokine in HCAEC. In normal conditions, GM CSF regulates alveolar clearance; however, during severe infections, it recruits inflammatory cytokines [20]. Overproduction of GM-CSF is related to harmful hyperinflammatory responses to COVID-19 [21,22] and can lead to endothelial dysfunction [23]. However, the changes in GM-CSF secretion were not observed in HLMVEC and HBEC cell cultures. Moreover, there were no changes in the level of other investigated cytokines after SCoV2-RBD treatment, and in HBEC cultures, the level of the secreted cytokines was extremely low. The fact we did not observe the concentration change after 24 hours of treatment with SCoV2-RBD might be due to an assessment timing mismatch with the cytokine secretion peaks. Additionally, cytokine synthesis and their secretion might be controlled by different mechanisms, the expression being related to preparation for the immune response, and the secretion – participating in the immune response after a particular trigger.

Summarising, the study reveals that SCoV-2-RBD triggers inflammatory gene upregulation in endothelial and epithelial cells primarily through the mitoROS – GM-CSF pathway, which in HLMVEC cells is also accompanied by acute mitochondrial network, cristae structure and function changes. The findings suggest mitoROS scavengers and/or antioxidants as potential therapeutics for preventing severe complications during coronavirus infections.

## 4. Materials and Methods

### 2.1 Cell culture and treatments

Primary human lung microvascular and coronary artery endothelial cells (HLMVEC and HCAEC, Cell Applications), and hTERT-immortalised bronchial epithelial cells (HBEC, Evercyte) were grown in Microvascular Endothelial Cell Growth Medium (Cell Applications), MesoEndo Cell Growth Medium (Cell Applications) and MEMα (Gibco, Life Technologies), respectively, supplemented with 10% FBS and 1% penicillin/streptomycin. MEMα for HBEC also contained 5 ng/mL EGF and 1 μg/mL hydrocortisone. Cells were cultured until 70-80% confluency, plated for experiments, and 24 hours later treated with SARS-CoV 2 Spike glycoprotein receptor binding domain protein (ScoV2-RBD, 2.8 μg/mL; Baltymas) with and without a specific mitochondrial superoxide scavenger MitoTempo (10 μM; Sigma-Aldrich) for 24 hours.

### 2.2 Mitochondrial morphology

6 × 10^4^ cells were seeded onto the glass part of P35 confocal dishes (75856-742, VWR), treated, stained with 100nM Mito Live Orange (Abberior) for 40 min and imaged using a fluorescent microscope Zeiss Axio Observer.Z1. Mitochondrial morphology was analysed using the Fiji MiNa toolset (Fig. A1). Images were preprocessed using unsharp masking, CLAHE, and median filtering. MiNa plugin was used to calculate the area of mitochondrial structures and produce a morphological skeleton for calculating parameters: Mean branch length, Summed branch length mean, and Mean of network branches [24]. At least 10 cells per experimental group were examined. Cristae were observed by an Olympus IX83 microscope with a STEDYCON STED nanoscope, UPLXAPO100xO NA1.45 objective. Parameters: pixel sizes 10-20, 2-5 line steps, dwell times 10 μs, pinhole 40 μm. Images were deconvoluted using Huygens software. Distances between cristae were measured using fluorescence intensity profiles along 1-2 μm line segments. Statistical analysis was conducted on the averages from 14-16 images.

### 2.3 Mitochondrial and glycolytic activity

Cells were seeded into the Seahorse XFp well plates at a density of 6 × 10^3^ cells/well. Mitochondrial and glycolytic function was assessed by the Seahorse XFp analyser, Cell Mito Stress Test Kit (Agilent Technologies). Final inhibitor concentrations in the wells were 1.5 μM oligomycin, 0.5 μM FCCP, 0.5 μM antimycin A, 0.5 μM rotenone. Oxygen consumption rate (OCR) and extracellular acidification rate (ECAR) were normalised to total protein content determined by Bradford assay. Optical density after reaction with the Bradford reagent (Merck) was assessed by Varioskan™ LUX Multimode Microplate Reader (Thermo Fisher Scientific Inc., Waltham, MA, USA). The data were analysed by Wave software.

### 2.4 Mitochondrial superoxide

Cells were seeded in 96-well plates at 104 cells/well. After treatments, the cells were stained with 2 μM MitoSox Red (Thermo Fisher Scientific) for 30 min in HBSS at 37 °C and imaged with an Olympus fluorescent microscope APX100. Mitochondrial location of MitoSOX Red was confirmed by co-staining with 500 nM MitoTracker Green (Thermo Fisher Scientific, Fig. A2a). Antimycin A (50 μM) was used 30 min prior to the cell staining as a positive control, and MitoTempo (10 μM, Sigma-Aldrich) as a negative control. Fluorescence intensity of the cells was measured using Olympus cellSens software selecting cell area segmentation from the background function (Fig.2b). Each measurement was performed in triplicate, with at least 10 images taken per replicate.

### 2.5 Gene expression

Cells were seeded in 12-well plates at 10^5^ cells/well, treated, total RNA extracted from the cells using PureLink RNA Mini Kit and reverse-transcribed by the High-Capacity cDNA Reverse Transcription Kit. Real-time quantitative PCR with Power SYBR Green Chemistry was used to evaluate the expression of IL-6, IL-8, IL-1β, NF-kβ, GM-CSF and CCL2 genes. The GAPDH gene was used as endogenous control. Sequences of PCR primers used in gene expression evaluation are presented in Table B1. Gene expression changes were normalised to untreated cells and represented as fold changes. All reagents were from Thermo Fisher Scientific.

## 5. Conclusions

- In human lung microvascular endothelial cells, SARS-CoV2-RBD fragments mitochondrial network, decreases cristae density and inhibits mitochondrial and glycolytic activity. The effects are not observed in coronary artery endothelial or bronchial epithelial cells.
- SARS-CoV2-RBD significantly increases mitochondrial superoxide in lung microvascular and coronary artery endothelial cells.
- Mitochondrial superoxide mediates SARS-CoV-2 RBD-induced expression of GM-CSF and IL-1β in lung and cardiac endothelial and bronchial epithelial, but IL-8 - only in the endothelial cells.

## Author Contributions

Conceptualization, A.J. and E.S.; methodology, G.K., D.N., Z.B., and D.G.; formal analysis, G.K., D.N., Z.B., D.G., M.K., and M.U.; investigation, G.K., D.N., A.T., A.V., Z.B., O.K., D.G., M.K., and M.U.; resources, A.J., Z.B., E.S.; writing—original draft preparation, D.N., G.K., A.J.; writing—review and editing, D.N., G.K., A.J., E.S.; visualisation, G.K., D.N., D.G., M.K., M.U., and A.J.; supervision, Z.B., A.J., E.S.; project administration, A.J., E.S.; funding acquisition, A.J., E.S. All authors have read and agreed to the published version of the manuscript.

## Data Availability Statement

Data are available upon reasonable request to the corresponding author.

## Appendix A

**Figure A1.**
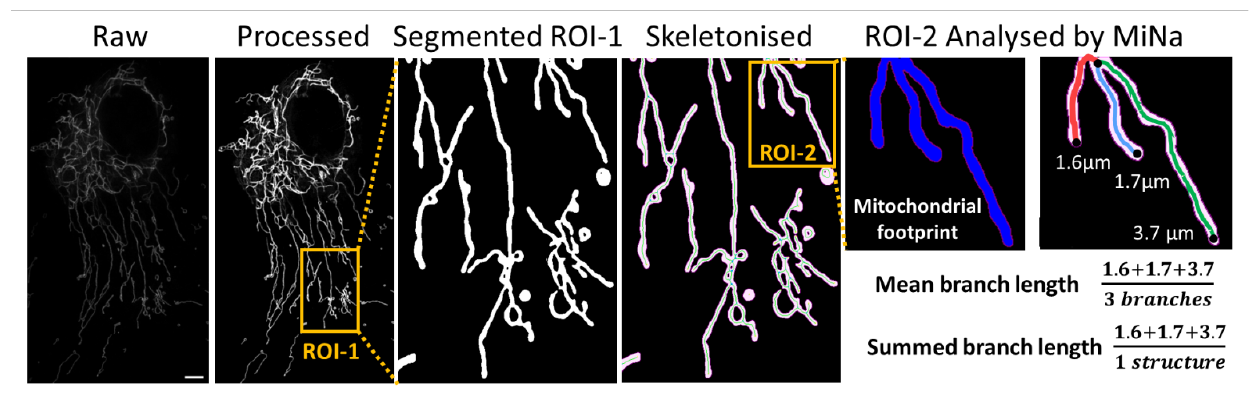
Mitochondrial network analysis steps and explanation of parameters. A representative STED microscopy image of a live untreated (control) human lung microvascular endothelial cell (HLMVEC) after Live Mito Orange staining. The scale bar is 2 μm. ROI – region of interest.

**Figure A2.**
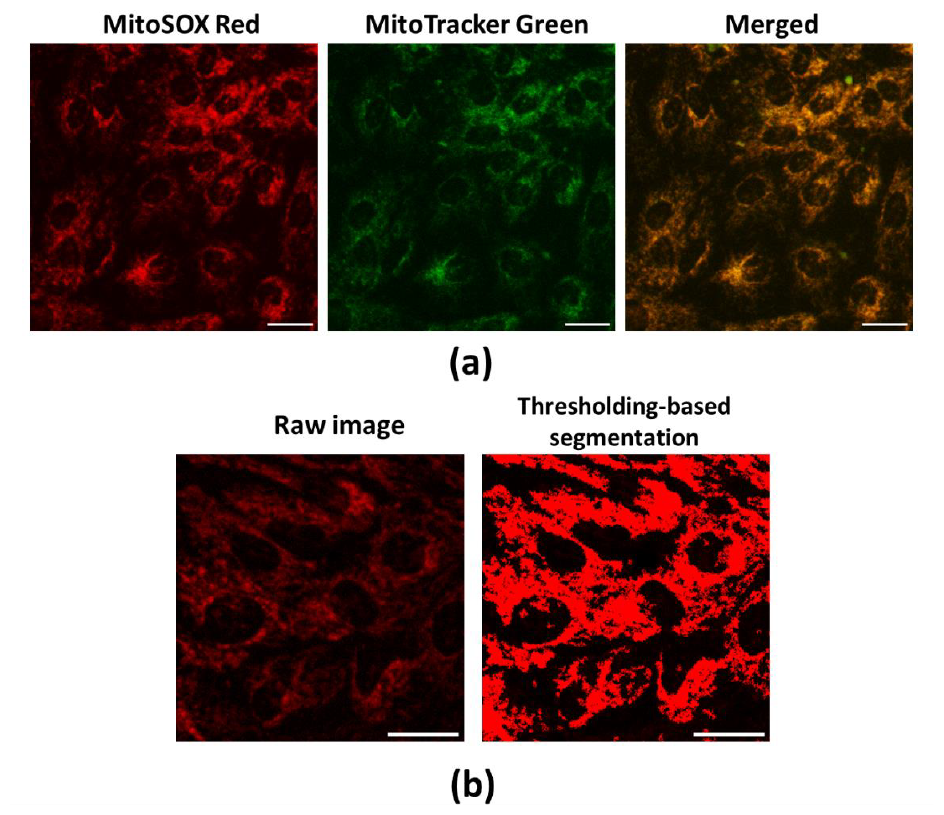
Representative images of mitoROS evaluation assay by mitoSOX. In (**a**) – an image representing mitoSOX Red colocalisation with mitochondrial network visualised by MitoTracker Green (a sample of SCoV2-RBD-treated human bronchial epithelial cells, or HBEC). The scale bar is 30 μm. In (**b**) – MitoSOX fluorescence intensity is evaluated by Olympus cellSens software in the area covered by cells, which is defined by the thresholding segmentation step (a representative image of HBEC control). The scale bar is 30 μm.

**Figure A3.**
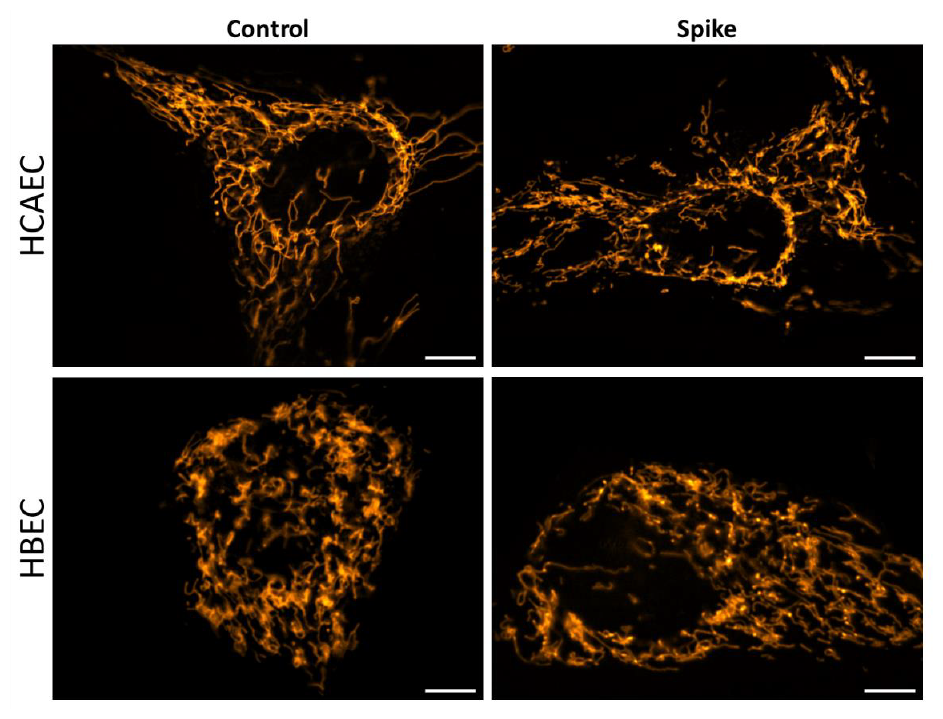
Representative images of mitochondrial morphology in HCAEC and HBEC after treatment with SCoV2-RBD (Spike). The scale bar is 10 μm.

**Figure A4.**
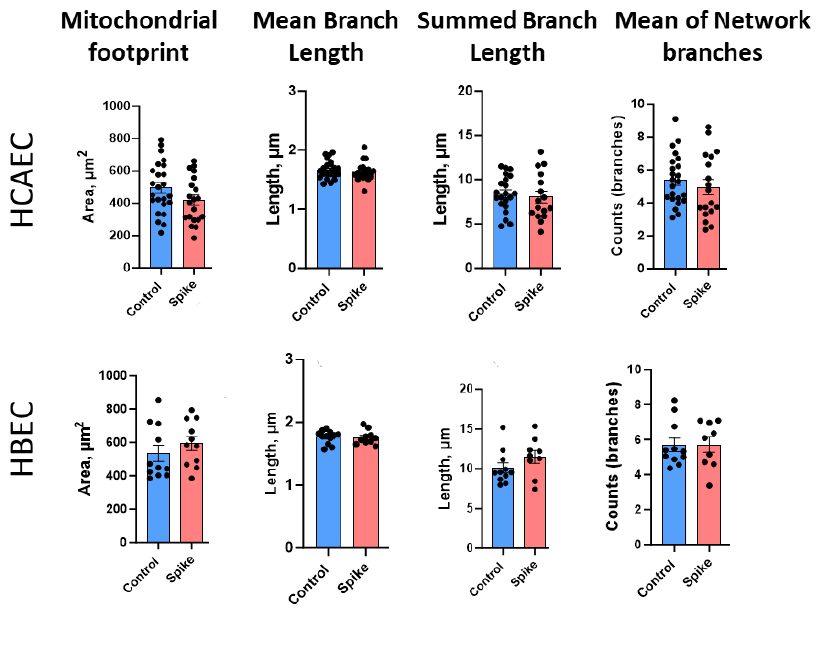
The quantitative results of mitochondrial network parameters of HCAEC and HBEC after treatment with SCoV2-RBD (Spike) calculated by means of the MiNa plugin of Fiji software.

**Figure A5.**
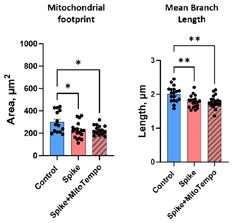
The impact of mitoROS in SCoV2-RBD-induced mitochondrial network alteration in HMLVEC. Spike - SCoV2-RBD. Results are normalised to untreated cells (Control) and represented as fold changes. ^*^p<0.05, ^**^p < 0.01; one way ANOVA Tukey’s test.

**Figure A6.**
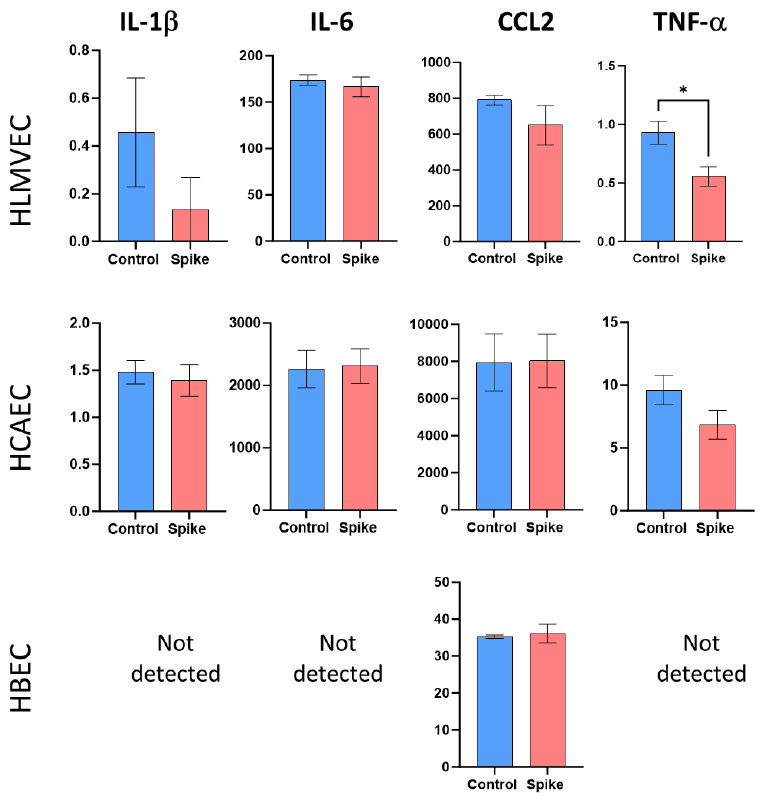
Proinflammatory cytokine secretion levels in HLMVEC, HCAEC and HBEC culture medium after SCoV2-RBD (Spike) treatment. *p<0.05; Student’s t-test.

## Appendix B

**Table B1.**
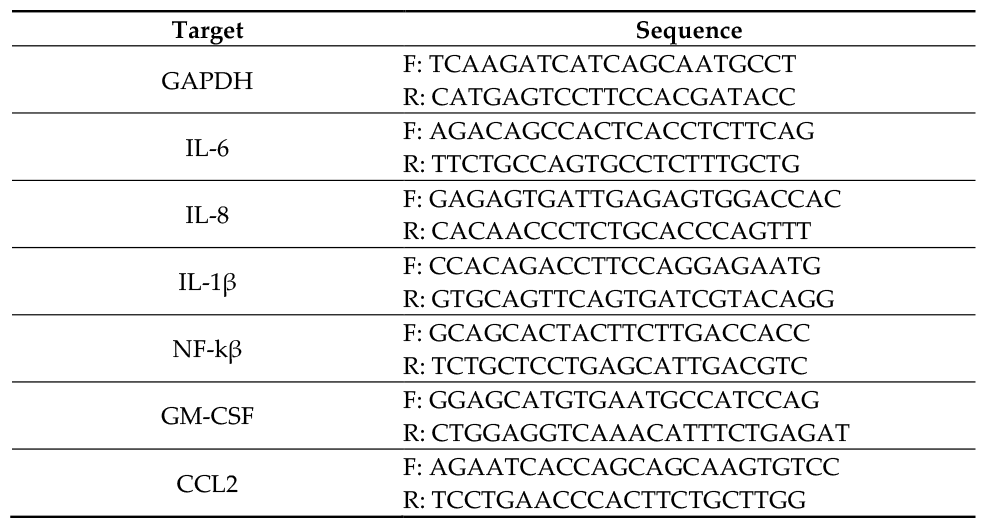
Sequences of PCR primers used for gene expression assessment.

---

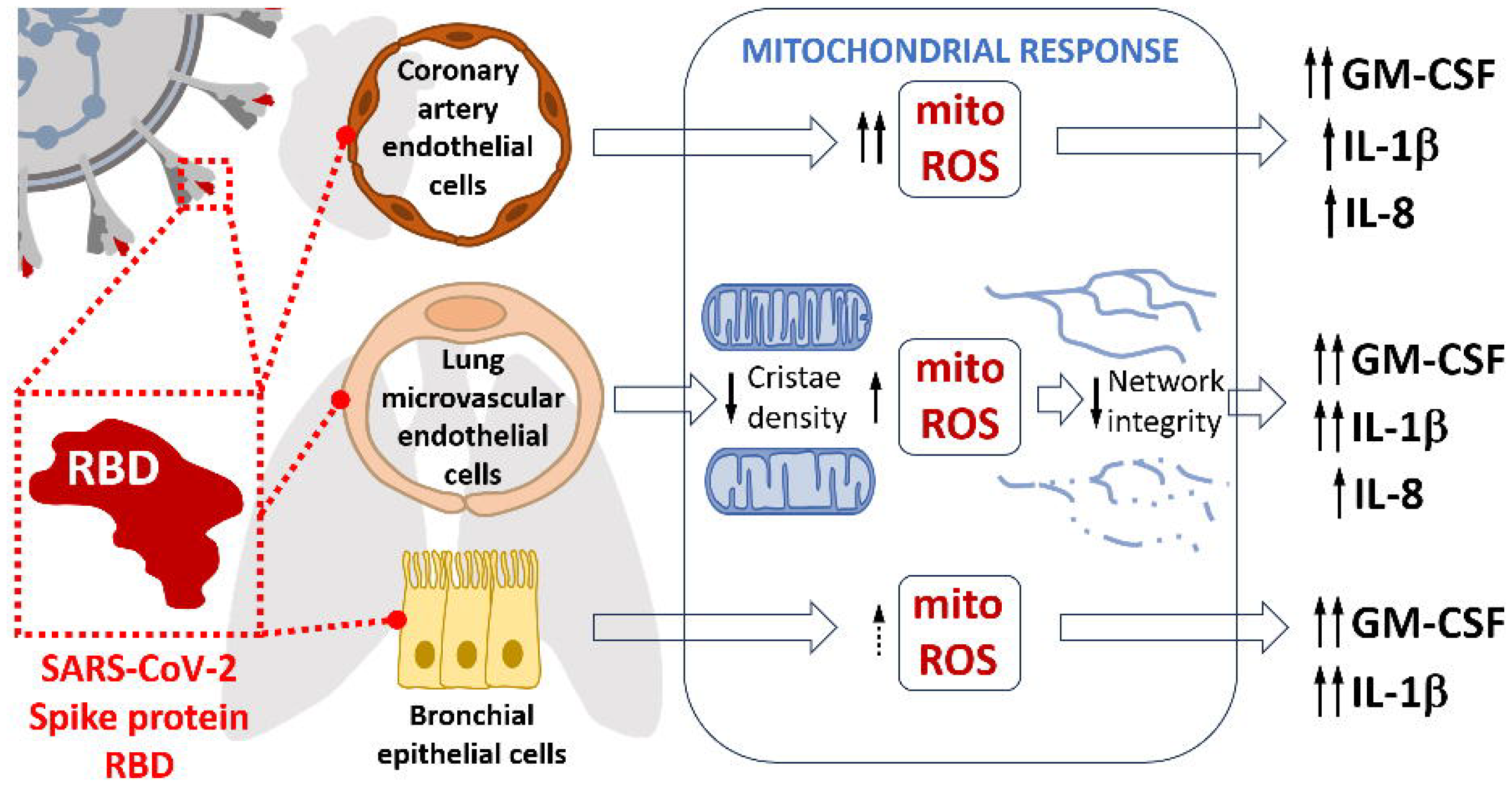

